# Sustaining attention for a prolonged period of time increases temporal variability in cortical responses

**DOI:** 10.1101/501544

**Authors:** Leon C. Reteig, Ruud L. van den Brink, Sam Prinssen, Michael X Cohen, Heleen A. Slagter

## Abstract

Our ability to stay focused is limited: prolonged performance of a task typically results in mental fatigue and decrements in performance over time. This so-called vigilance decrement has been attributed to depletion of attentional resources, though other factors such as reductions in motivation likely also play a role. In this study, we examined three EEG markers of attentional control, to elucidate which stage of attentional processing is most affected by time-on-task and motivation. To elicit the vigilance decrement, participants performed a sustained attention task for 80 minutes without breaks. After 60 minutes, participants were motivated by an unexpected monetary incentive to increase performance in the final 20 minutes. We found that task performance and self-reported motivation declined rapidly, reaching a stable levels well before the motivation manipulation was introduced. Thereafter, motivation increased back up to the initial level, and remained there for the final 20 minutes. While task performance also increased, it did not return to the initial level, and fell to the lowest level overall during the final 10 minutes. This pattern of performance changes was mirrored by the trial-to-trial consistency of the phase of theta (3–7 Hz) oscillations, an index of the variability in timing of the neural response to the stimulus. As task performance decreased, temporal variability increased, suggesting that attentional stability is crucial for sustained attention performance. The effects of attention on our two other EEG measures—early P1/N1 event-related potentials and pre-stimulus alpha (9–14 Hz) power—did not change with time-on-task or motivation. In sum, these findings show that the vigilance decrement is accompanied by a decline in only some facets of attentional control, which cannot be fully brought back online by increases in motivation. The vigilance decrement might thus not occur due to a single cause, but is likely multifactorial in origin.

## 1. Introduction

Our ability to stay focused is not limitless. When a task requires sustained attention for a prolonged period of time, performance on that task typically decreases, and mental fatigue increases (Ackerman, 2011). This phenomenon is known as the time-on-task effect or the vigilance decrement (Warm, Parasuraman, & Matthews, 2008). Unfortunately, the vigilance decrement is most pronounced in situations where its consequences can be dire: in airport security personnel looking for suspicious objects in luggage, lifeguards scanning the horizon for potential drowning victims, or truck drivers that spend long periods of time on the road. What these scenarios have in common is a high rate of stimuli that require no action (e.g., trees next to the road), coupled with signals that are rare and easy to miss, but critical (e.g., a dog about to cross the road) (Parasuraman, 1979). These principles have been translated into more simple experimental tasks, in which case the vigilance decrement is also observed (Bartlett, 1943; Mackworth, 1948).

From decades of research, three broad classes of frameworks have emerged that attempt to explain the origin of the vigilance decrement in terms of *overload, underload*, and *motivational control. Overload* theories mainly attribute the vigilance decrement to depletion of cognitive resources (Helton & Warm, 2008). In this view, successful monitoring of incoming information requires allocation of limited resources to the task at hand. When vigilance has to be maintained over time, the pool of resources is steadily depleted, causing task performance to progressively worsen.

Alternatively, *underload* accounts hold that typical vigilance tasks are simply not stimulating enough to continue to capture attention, causing people to disengage from the task (Manly, Robertson, & Galloway, 1999; Robertson, Manly, Andrade, Baddeley, & Yiend, 1997). As attention is directed away from the task, participants also become occupied with task-unrelated thoughts—they start mind wandering (Smallwood & Schooler, 2006), and this is what causes performance to drop.

Yet other factors, such as motivation, likely also play an important role. *Motivational control theories* (Hockey, 1997) are based on the premise that sustained attention requires sufficient motivation and low levels of aversion to the task at hand. Task performance is only maintained if a cost-benefit analysis shows that the rewards obtained by doing so (e.g., appreciation from others) offset the costs (e.g., stress). Feelings of fatigue and effort expenditure are experienced when instead the costs start to outweigh the benefits (Kurzban, Duckworth, Kable, & Myers, 2013).

Some more recent proposals recognize that these three classes are not mutually exclusive. For instance, the resource control model synthesizes overload and underload accounts, by explaining the vigilance decrement as a disproportionate amount of resources that are occupied by mind wandering (Thomson, Besner, & Smilek, 2015). Likewise, the beneficial effects of motivation on task performance may be mediated by mind wandering (Mrazek et al., 2012) as more motivated participants tend to have less task-unrelated thoughts (Seli, Cheyne, Xu, Purdon, & Smilek, 2015; Seli, Schacter, Risko, & Smilek, 2017). Other hybrid models have combined the resource depletion and motivational control accounts by construing cost-benefit analysis mainly in terms of energy expenditure (Boksem & Tops, 2008; Christie & Schrater, 2015).

All in all, it is not yet fully understood why it is so difficult to sustain attention for a prolonged period of time. A better understanding of the neural mechanisms underlying the vigilance decrement may help to answer this question. A large body of research has shown— using the high-temporal resolution of EEG—that attention can change neural processing at different stages. Tracking these neural markers of attention over time may elucidate how time-on-task changes information processing.

First, the ability to sustain attention (as indexed by reaction time variability) has been linked to neural response variability, specifically the cross-trial consistency of the phase of post-stimulus theta oscillations (Lutz et al., 2009). However, no studies have examined how this theta phase response changes with time-on-task. Second, attention can enhance early visual processing, as reflected in the amplitude of the visual-evoked P1 and N1 components (Eason, Harter, & White, 1969; Luck et al., 1994; Mangun & Hillyard, 1991). But it remains unclear how P1 and N1 amplitude are affected by time-on-task: some studies have demonstrated decreases in N1 amplitude with time-on-task (Boksem, Meijman, & Lorist, 2005; Faber, Maurits, & Lorist, 2012), but others report that the N1 remains stable (Bonnefond, Doignon-Camus, Touzalin-Chretien, & Dufour, 2010; Koelega et al., 1992). Finally, top-down attention can bias visual regions in advance through modulations of pre-stimulus alpha-band activity (Mazaheri et al., 2014; O’Connell et al., 2009). Although some studies have shown that alpha power increases with time-on-task (Boksem et al., 2005; Bonnefond, Doignon-Camus, Hoeft, & Dufour, 2011; Bonnefond et al., 2010), this could also reflect changes in general arousal or alertness (Cajochen, Brunner, Krauchi, Graw, & Wirz-Justice, 1995; Drapeau & Carrier, 2004), rather than a specific change in top-down attention.

In the current study, we used these three EEG markers to track changes in attention with time-on-task. Each indexes a different stage of attentional processing: late post-stimulus (theta phase), early post-stimulus (P1/N1 components) and pre-stimulus (alpha power). Our approach might thus yield a better understanding of which specific attentional processes deteriorate over time with prolonged sustained attention performance. To this end, we designed an experiment that allowed us to examine changes in attentional control itself, teasing it apart from other factors like changes in general arousal or motivation. EEG was continuously recorded while participants performed a sustained attention task for 80 minutes (without breaks) in which they had to detect rare targets. We expected to find the classic vigilance decrement, i.e. that participants’ ability to discriminate targets from non-targets would decrease with time-on-task.

We also aimed to explore how changes in motivation would interact with the vigilance decrement and its neural correlates. To do so, after 60 minutes of time-on-task, we motivated participants to increase their performance by offering the prospect of a monetary reward that was contingent on adequate task performance. Several studies have shown that social comparison or monetary rewards can improve performance (Boksem, Meijman, & Lorist, 2006; Bonnefond et al., 2011; Hopstaken, van Der Linden, Bakker, & Kompier, 2015; Lorist et al., 2009). Yet, it is still unclear whether motivation acts directly upon neural mechanisms that are critical to sustaining attention, or if motivation enhances task performance through other means.

We formulated specific predictions for each of our three EEG measures. First, we hypothesized that the EEG response evoked by the stimulus at later stages of information processing would become less stable over time, as attention becomes more reactive with increasing time-on-task. The consistency in the timing of stimulus-evoked responses can be quantified with inter-trial phase clustering (ITPC), a measure based on the phase angles of oscillatory neural activity (VanRullen, Busch, Drewes, & Dubois, 2011). ITPC in the theta (4–8 Hz) band was reported to increase following meditation training (Lutz et al., 2009; Slagter, Lutz, Greischar, Nieuwenhuis, & Davidson, 2009), suggesting that training enhanced attentional stability. Here, we expect to find the opposite: because attentional stability should degrade with time-on-task, resulting in more variability in the timing of stimulus processing, theta ITPC should decrease. Although to our knowledge no studies have specifically examined time-on-task effects, earlier studies have reported that fatigue (resulting from sleep deprivation) may decrease theta ITPC (Eidelman-Rothman et al., 2018; Hoedlmoser et al., 2011).

Second, if time-on-task affects early visual processing, the effect of top-down attention on P1/N1 amplitude should weaken over time. That is, the difference between trials in which attention was successfully deployed (resulting in a correct identification of the target—a hit) and trials in which attention lapsed (resulting in a miss) should decrease. Two earlier studies reported decreases in absolute N1 amplitude over time in spatial cueing (Boksem et al., 2005)and flanker (Faber et al., 2012) tasks, but no change in P1 amplitude. However, these reductions in absolute N1 amplitude could also reflect habituation. Furthermore, two other studies using more traditional sustained attention tasks found that N1 amplitude remained stable with time-on-task (Bonnefond et al., 2010; Koelega et al., 1992).

Finally, time-on-task might also degrade preparatory orienting of attention, as indexed by oscillatory activity in the alpha band (8–14 Hz). A large body of work has demonstrated that spatial attention can be read out by examining the topographical pattern of alpha power prior to stimulus presentation. For example, when attention is directed to one hemifield (e.g., left) in expectation of a relevant stimulus, alpha power over the ipsilateral (left) hemisphere increases, while alpha power over the contralateral (right) hemisphere decreases (Sauseng et al., 2005; Thut, Nietzel, Brandt, & Pascual-Leone, 2006; Worden, Foxe, Wang, & Simpson, 2000). We thus designed our task such that stimuli were only presented left of fixation, requiring participants to covertly deploy attention to one visual hemifield. We expected the typical asymmetry in alpha power to decline with time-on-task, reflecting a decreased ability to direct attention in preparation of the stimulus. Such top-down attention related-changes can be separated from more general, alertness-related increases in alpha power, which do not have this hemisphere-specific signature (Cajochen et al., 1995; Drapeau & Carrier, 2004). One earlier study showed that alpha power becomes more right-lateralized with time-on-task (Newman, O’Connell, & Bellgrove, 2013), but because stimulus location was not cued in this study, this lateralization likely did not reflect a change in top-down orienting of attention.

## 2. Materials and Methods

Following Simmons, Nelson, & Simonsohn (2012), we report how we determined our sample size, all data exclusions, all data inclusion/exclusion criteria, whether inclusion/exclusion criteria were established prior to data analysis, all manipulations, and all measures in the study. This study was not preregistered.

A different set of results based on this dataset has been reported in an earlier publication (Slagter, Prinssen, Reteig, & Mazaheri, 2016). The experimental task, procedure and participant population is identical.

### 2.1. Participants

Data of 21 participants (11 female, mean age = 21.6, *SD* = 3.4, range = 18–26) are reported here. A total of 30 people participated in the study, but nine were excluded from data analysis due to a malfunctioning reference electrode (5 participants), poor EEG data quality (2 participants,) not completing the task (1 participant), or problems maintaining fixation (1 participant). Our sample size of 30 participants was determined a priori, by comparison to previous studies of the time-on-task effect (MacLean et al. (2009): n =17; Lorist et al. (2009): n = 15, Boksem et al. (2005): n = 17), as well as the literature on attentional modulations of the P1/N1 (e.g. Talsma, Mulckhuyse, Slagter, & Theeuwes (2007): n = 16; Grent-’t-Jong & Woldorff (2007): n = 13). The precise exclusion criteria were not determined a priori, but we did anticipate having to exclude some participants due to EEG data quality issues, hence we included more participants than the final sample size of most earlier studies. Participation was precluded in case someone reported getting at least 2 hours less sleep than usual the night before the experiment. The experiment and recruitment took place at the University of Amsterdam. All procedures for this study were approved by the ethics review board of the Faculty for Social and Behavioral Sciences. All participants provided their written informed consent and were compensated with course credit or €7 per hour. A subset (35%) of participants also received another €30 based on their task performance (see Procedure).

### 2.2. Task

Participants performed a modified version of the sustained attention task from MacLean et al. (2009) (exp. 1, stable version), run using Presentation (Neurobehavioral systems, Inc.). This task required participants to respond when they detected a rare target signal (short lines), but to withhold a response for standard non-targets (long lines) (Figure 1).

**Figure 1:**
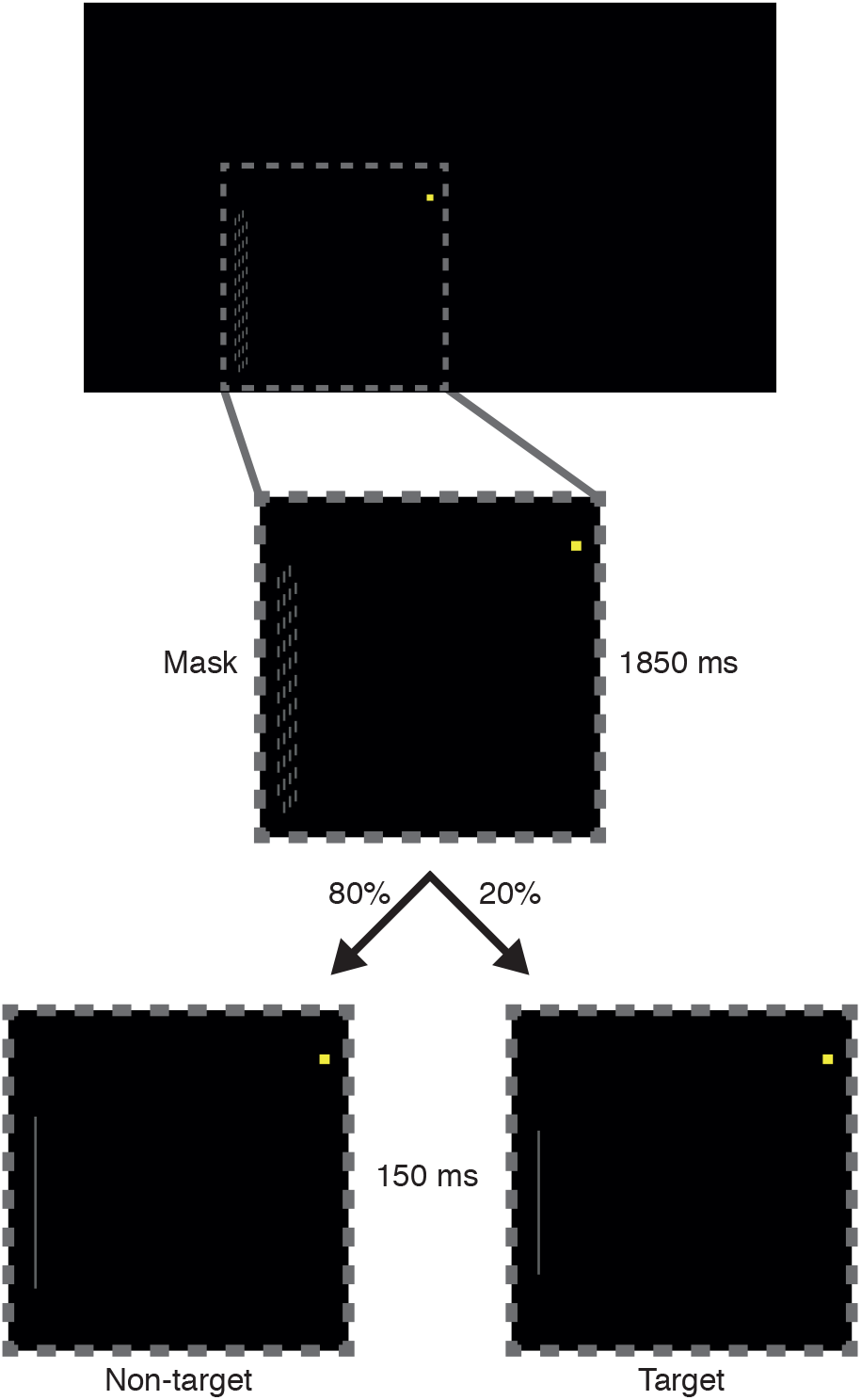
Sustained attention task. All stimuli were presented 3° to the left and 1.5° below the yellow fixation square. The inset (dotted square) depicts a zoomed view of the stimuli. Each trial started with the presentation of the mask stimulus for 1850 ms. Then either a non-target (long line; 80% of trials) or a target (short line; 20% of trials) was presented for 150 ms. Participants were instructed to only respond to targets.

Stimuli were presented at a viewing distance of 110 cm against a black background on a 17-inch Benq TFT monitor, running at a 60 Hz refresh rate. Participants were instructed to maintain fixation on a central yellow square (0.11° x 0.11°), which remained on screen throughout the task. All other stimuli were presented 3° to the left and 1.5° lower than fixation, so participants had to continuously and covertly direct their attention to this location. Every 2000 ms, a light gray line (width: 0.03°) was presented for 150 ms, which could be either short (a target; 20% of trials) or long (a non-target; 80% of trials). Participants were tasked to respond with their right index finger whenever they detected a target (short line). Non-targets were always 1.89° long; target length was calibrated individually (see Procedure) for each participant (*M* = 1.40°, *SD* = 0.10°, range = 1.21–1.59°). Line stimuli were preceded and followed by a mask stimulus, presented for the remaining 1850 ms. The mask was composed of many short lines (0.03° x 0.12°), subtending an area of 0.21° by 2.44°. To prevent participants from judging the length of (non-)target lines relative to the mask, the lines that comprised the mask were vertically shifted by a random small amount (within ±0.06°) on each presentation.

### 2.3. Procedure

Before the main task, the level at which individual participants performed the task was titrated to achieve a starting point of 80% accuracy (i.e. the target was detected 80% of the time). We did so to equate the task demand across participants and to maximize sensitivity to changes in performance with time-on-task. As in MacLean et al. (2009), task difficulty was adjusted by calibrating the length of the target line using Parameter Estimation by Sequential Testing (PEST) (Taylor & Creelman, 1967). PEST is an adaptive thresholding procedure in which the stimulus is adjusted until a stable level of performance is reached. During the PEST procedure, participants received auditory feedback for incorrect (misses and false alarms) or correct (hits only) responses (a “ding” sound for correct; a “whoosh” sound for incorrect). The only other difference between the PEST- and main task versions was a higher target rate (PEST: 1 in every 3.5 trials; main task: 1 in 5), to more quickly estimate the threshold target length.

The time it took to complete the PEST procedure varied modestly between participants (range: 7–13 minutes). Afterwards, participants performed 2400 trials (480 target trials) of the main task, which lasted for 80 minutes. No breaks were included, to prevent participants from mitigating the vigilance decrement by taking rest.

There were only two different interruptions of the task. First, before starting the task and every 10 minutes thereafter (after 300 trials), participants rated their levels of motivation (“Please rate your motivation for performing well on this task”) and aversion (“Please rate your aversion towards this task”) on a 7-point scale (1: “not motivated”/”no aversion”, 7: “highly motivated”/“strong aversion”). Participants had 6 seconds do to so per rating, by moving a cursor along the scale to the number of their choosing.

Second, after 60 minutes, a new screen was displayed informing participants of the chance to gain an additional monetary reward—an option that was unbeknownst to them until then. This manipulation was designed to evaluate whether motivation could counteract the decline in vigilance over time. Participants could gain €30 on top of their normal compensation, if they outperformed at least 65% of the other participants during the final 20 minutes of the task (cf. Lorist et al. (2009); see the Appendix for the full instruction text). These instructions were presented for a maximum of 60 seconds, or until the participant indicated they read them by pressing the mouse button. All participants pressed within 60 seconds, after which they rated their motivation and aversion one additional time (such that we could compare these ratings immediately before and after the motivation manipulation) before the task continued.

### 2.4. Behavioral data analysis

We computed *A*′, a non-parametric index of perceptual sensitivity (Stanislaw & Todorov, 1999), to assess participants’ ability to discriminate targets from non-targets (cf. MacLean et al. (2009)):

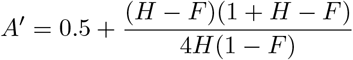

if *H* > *F*, where *H* is hit rate (*Hits/(Hits + Misses*)) and *F* is false alarm rate (*False alarms/(False alarms* + *Correct rejections*)). *A*′ can take any value between 0.5 (targets are indistinguishable from non-targets) and 1 (perfect discriminability). To track changes in performance over time, *A*′ was computed for each block of 10 minutes of the experiment (8 blocks in total).

We also analyzed mean response times in hit trials per 10-minute block. Initially, we also planned to analyze the response time coefficient of variation (mean divided by standard deviation), which is known to increase with time-on-task (Esterman, Noonan, Rosenberg, & Degutis, 2013; van den Brink, Murphy, & Nieuwenhuis, 2016). However, precisely estimating the variability in response time requires more trials with responses than we were left with (Hit trials per 10-minute block: *M* = 33, *SD* = 10.5, range = 10–54).

### 2.5. EEG data analysis

#### 2.5.1. Acquisition and preprocessing

EEG data were recorded at 512 Hz using a BioSemi ActiveTwo system with 64 Ag/AgCl active electrodes, placed according to the (10-10 subdivision of the) international 10-20 system. Two pairs of additional external electrodes were placed to record the EOG, above and below the left eye, and next to the left and right outer canthi.

Preprocessing was performed using the EEGLAB toolbox (Delorme & Makeig, 2004) in MATLAB (Mathworks, Inc.). The continuous EEG data were high-pass filtered at 0.1 Hz using a Hamming-windowed sinc FIR filter. Data were then segmented into epochs from −2000 ms to 3000 ms peri-stimulus (non-targets or targets), including buffer zones on either end to accommodate the edge artifacts that may result from wavelet convolution (see Time-frequency decomposition). Epochs containing muscle activity, eye movements or large artifacts due to other sources of noise were removed after visual inspection. Bad channels were interpolated with spherical spline interpolation. Subsequently, independent component analysis was performed on all channels (including the EOG channels). Components consisting of eye blinks or other artifacts clearly distinguishable from neural activity were subtracted from the data. Finally, epochs were average referenced and separated into correct rejections, hits and misses. False alarm trials were too few (only 3% of non-target trials) to include in the EEG analysis.

#### 2.5.2. Trial binning

Corresponding to the behavioral data, correct rejection trials were binned into eight blocks of 10 minutes of task performance. We equated trial numbers across blocks for each participant, by discarding a (randomly selected) subset of trials from blocks that had more trials than the minimum. This was done because our inter-trial phase clustering measure (see Time-frequency decomposition) in particular can be biased by differences in trial counts between conditions (Cohen, 2014). After this subsampling procedure, an average of 167 correct rejection trials (*SD* = 22.5, range = 124–213) remained per participant for each block. 20-minute bins (4 blocks in total) were used for hit and miss trials, as these were less frequent. Hit and miss trials were also subsampled such that the same amount of hit and miss trials were analyzed per block (*M* = 24, *SD* = 5.4, range = 11–33).

This subsampling procedure was repeated 1000 times; each time a trial-average of each measure (see Event-related potentials, Time-frequency decomposition) was computed. All 1000 subsampled trial-averages were then averaged together, such that the final value should reflect the average across all trials. The subsampling was repeated in this manner to prevent biases in trial selection, for example because the subsample happened to contain mostly trials from the first half of a block.

#### 2.5.3. Event-related potentials

For the ERP analysis, epochs were baseline corrected from −200 to 0 ms pre-stimulus, and averaged separately for each condition. The resulting ERPs were low-pass filtered at a cut-off of 30 Hz (for plotting purposes, for statistical analysis no low-pass filter was applied).

#### 2.5.4. Time-frequency decomposition

Time-frequency representations of the EEG data were obtained through complex Morlet wavelet convolution (Cohen, 2014). Wavelet frequency increased from 2 to 80 Hz in 30 logarithmically spaced steps. The number of wavelet cycles was increased from 3 to 12 in the same number of steps, to increase temporal precision at lower frequencies and frequency precision at higher frequencies.

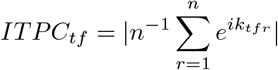

where *n* is the number of trials, and *e^ik^* is the complex representation of phase angle *k* on trial *r* at time-frequency point *tf. ITPC* is low when phase values are uniformly distributed across trials (bounded at 0, meaning the phase angle at a certain time is completely random from trial to trial). *ITPC* is high when phase angles cluster around a preferred value across trials (bounded at 1, meaning the phase angle at a certain time is exactly the same on each trial).

We also extracted power at each time point and frequency, which was subsequently baseline corrected using a decibel conversion (Cohen, 2014): 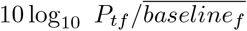, where *P_tf_* is power at time-frequency point *tf* and 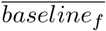 is power at frequency *f*, averaged across a −400 to −100 ms window relative to stimulus onset.

Finally, to examine spatial attentional bias, we calculated a lateralization index of trial-averaged power values (again per frequency and time point):

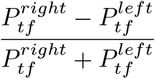

where 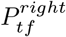 is power at time-frequency point *tf* at an electrode on the right side of the midline (e.g. C4) and 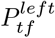 is the equivalent electrode on the left side (e.g.C3). The index is positive when 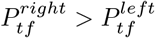, and negative for the inverse case. Because this analysis was aimed at examining pre-stimulus power, we did not apply a baseline correction, but instead normalized by total power 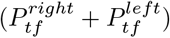 (cf. Händel, Haarmeier, & Jensen, 2011).

### 2.6. Statistical analysis

*A*′, hit rate, false alarm rate, and response time were analyzed separately using one-way repeated measures ANOVAs with the within-subject factor Block (eight 10-minute blocks of task performance). If the ANOVA revealed a significant effect, we conducted paired t-tests to examine differences between block 1 (first 10 minutes of task performance), block 6 (last block before the motivation manipulation, see Procedure), block 7 (first block after the motivation manipulation), and block 8 (last block of task performance). Specifically, we planned pairwise comparisons of block 1 vs. block 6, block 1 vs. block 7, block 6 vs. block 7, and block 1 vs. block 8. Bayesian analogues of these t-tests (with a standard Cauchy prior) (Rouder, Speckman, Sun, Morey, & Iverson, 2009) were used to quantify the relative evidence for an effect (BF_10_) or for the null hypothesis (BF_01_). Motivation and aversion ratings were subjected to a 1×10 repeated measures ANOVA, as there were two ratings in addition to the one after each block: before the task, and directly after the motivation manipulation. For the planned paired tests, these two ratings were used instead of block 1 and block 7.

Similar to the behavioral analyses, EEG data were also subjected to repeated measures ANOVAs with the factor Block. We also followed up each ANOVA with a planned paired test between block 6 and 7, to quantify the effect of motivation. For all EEG analyses, the electrodes, time windows, and frequency windows of interest were determined based on visual inspection of the grand average plots (averaged over participants and blocks) in correct rejection trials.

To examine changes in attentional modulation of early visual processing, our ERP analyses focused on the P1 and N1 components. We extracted average voltage values per participant in a window of 17.5 ms on either side of the P1 and N1 peak. For each participant, the P1 peak latency was defined as the sample with the maximum (positive) voltage within 110–180 ms post-stimulus; N1 peak latency was defined as the sample with the minimum (negative) voltage within 190–260 ms post-stimulus. These values were averaged for two pools of electrodes: left parieto-occipital (PO5, P5, P7) and right parieto-occipital (PO6, P6, P8). We ran separate repeated measures ANOVAs for the P1 and N1: an 8 (Block) x 2 (Hemisphere: left PO vs. right PO) ANOVA for correct rejection trials, and another 4 (Block) x 2 (Hemisphere) x 2 (Condition: hits vs. misses) ANOVA.

To examine changes in attentional stability, we analyzed ITPC values for the same left PO and right PO electrode pools, as well as a mid-frontal (MF) pool of electrodes (FCz, FC1, FC2). Similar to the ERP analysis, we extracted average values for statistics using participant-specific windows of 60 ms and 1 Hz on either side of the peak value for that participant, within a larger window of 150–500 ms and 3–7 Hz (consistent across participants). We again ran two repeated measures ANOVAs: an 8 (Block) x 3 (Region: left PO vs. right PO vs. MF) ANOVA for correct rejection trials, and another 4 (Block) x 3 (Region) x 2 (Condition: hits vs. misses) ANOVA. We also analyzed theta power, using the exact same electrodes, time-frequency windows, and ANOVA factors.

In an additional post hoc analysis, we quantified the relationship over time between ITPC and *A*’ using a multilevel growth model. Multilevel growth models are an extension of linear mixed models that can accommodate hierarchical data structures (see e.g. Kristjansson, Kircher, & Webb, 2007). A multilevel growth model is appropriate in our case, because changes over time in variables at one level (ITPC, *A*’) are nested within another level (participants). Specifically, we modeled ITPC per 10-minute block as a fixed effect and *A*’ per block as the outcome. To account for the correlation between repeated measures, we specified a first-order auto-regressive covariance structure between the 8 blocks, nested within participants.

Finally, to examine changes in preparatory attentional orienting, we extracted the average pre-stimulus (−1000 to −100 ms) power lateralization index values in the alpha range (9–14 Hz), again from the left PO and right PO electrode pools, and subjected these to 1×8 (Block) (correct rejections) and 4 (Block) x 2 (Condition: hits vs. misses) repeated measures ANOVAs.

All statistical analyses were conducted using several R (R Core Team, 2017) packages (Lawrence, 2016; Morey & Rouder, 2018; Pinheiro, Bates, & R-core, 2018; Robinson & Hayes, 2018; Wickham, 2017) from within RStudio (RStudio Team, 2016). Effect sizes are reported as (partial) eta-squared 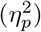 or Cohen’s *d. α* was fixed at 0.05. In case Mauchly’s test for violations of sphericity was significant, we report Greenhouse-Geisser corrected p-values and degrees of freedom.

### 2.7. Data, materials, and code availability

The data, materials, and analysis code can be obtained from this study’s page on the Open Science Framework^3^, including: The raw behavioral data, as well as the raw and preprocessed EEG data (Reteig, van den Brink, Prinssen, Cohen, & Slagter, 2018); An R notebook (Xie, 2015, 2018) that reproduces all of the statistical results in this publication; MATLAB scripts that reproduce all the figures in this publication; MATLAB code to compute the EEG measures from the preprocessed EEG data and extract the data for statistics.

## 3. Results

### 3.1. Behavior

The sustained attention task elicited a robust vigilance decrement: accuracy in discriminating targets from non-targets (defined as *A*′) decreased with time-on-task (Figure 2A, *F*(2.50, 49.96) = 7.47, *p* < .001, *η*^2^ = .27). Accuracy declined rapidly until around 30 minutes into the task, when it became more or less stable. Response times also followed this pattern, but stabilized earlier (Figure 2B, *F*(3.82, 76.45) = 2.56, *p* = .047, *η*^2^ = .11). With time-on-task, motivation to continue decreased (Figure 2E, *F*(3.89, 77.80) = 7.10, *p* < .001, *η*^2^ = .26), while aversion ratings increased (Figure 2F, *F*(3.46, 69.17) = 12.67, *p* < .001, *η*^2^ = .39).

**Figure 2:**
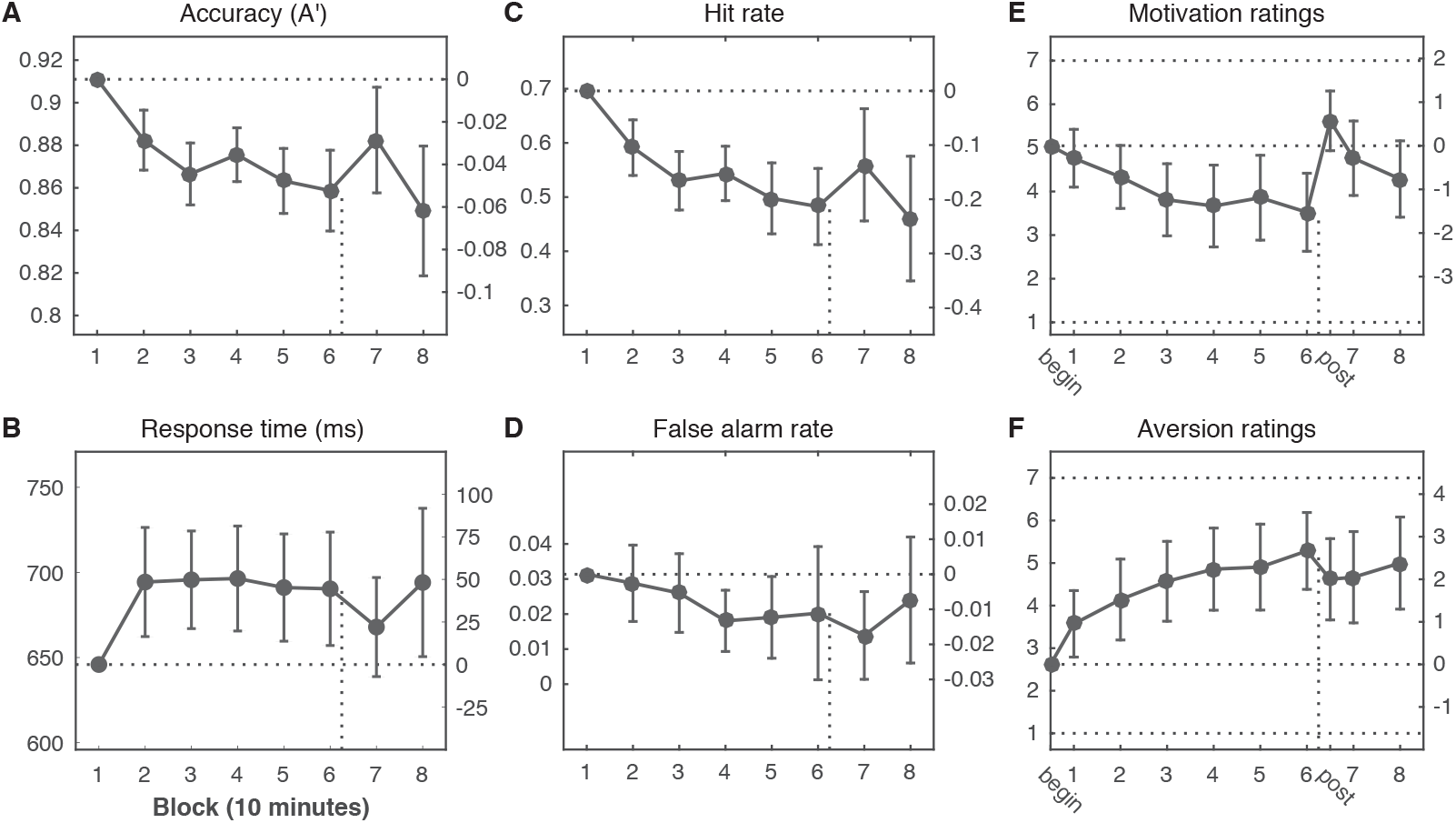
Declining performance with time-on-task is only partially counteracted by increases in motivation. (**A**) Accuracy (*A*′) declined with time on task and was only partially and transiently restored by the motivation manipulation (after 60 minutes of task performance; vertical dotted line). These changes in accuracy were mirrored in the response time data (**B**) and mostly due to changes in Hit rate (**C**); there was no significant change over time in False alarm rate (**D**). The motivation manipulation successfully restored motivation ratings to their initial levels (**E**), but aversion ratings (**F**) remained elevated. The left y-axis of each plot shows the absolute value; the right y-axis shows the change compared to the first measurement (horizontal dotted line). Error bars are 95% confidence intervals of the paired difference between the first and each subsequent mean; paired differences are significant when the confidence interval does not overlap the dotted line. The x-axis depicts time-on-task in blocks of 10 minutes. The motivation and aversion ratings (horizontal dotted lines: minimum of 1, maximum of 7) were taken after each block (1 through 8), as well as before the task started (*begin*) and directly after the motivation manipulation (*post*).

After one hour (after block 6), participants learned that they could earn an additional sum of money if they outperformed at least 65% of the other participants during the final 20 minutes of the task. Just before this manipulation, motivation was at its lowest (begin vs. 6: *M_diff_* = 1.52, *t*(20) = 3.55, *p* = .002, *d* = 0.78, BF_10_ = 19.9), and aversion was its peak (begin vs. 6: *M_diff_* = −2.67, *t*(20) = −6.16, *p* < .001, *d* = −1.34, BF_10_ = 4057). Right after the motivation manipulation, participants were significantly more motivated compared to before (6 vs. post: *M_diff_* = −2.10, *t*(20) = −5.14, *p* < .001, *d* = −1.12, BF_10_ = 515), and also displayed decreased aversion towards performing the task (6 vs. post: *M_diff_* = 0.67, *t*(20) = 3.16, *p* = .005, *d* = 0.69, BF_10_ = 9.2). Motivation ratings were no longer significantly different from the beginning (begin vs. post: *M_diff_* = −0.57, *t*(20) = −1.74, *p* = .097, *d* = −0.38, BF_01_ = 1.21), and remained at this level after the final two blocks (begin vs. 8: *M_diff_* = 0.76, *t*(20) = 1.82, *p* = .084, *d* = 0.40, BF_01_ = 1.09).

The motivation manipulation improved accuracy in the following block of task performance (6 vs. 7: *M_diff_* = −0.024, *t*(20) = −2.51, *p* = .021, *d* = −0.55, BF_10_ = 2.75). However, accuracy was still significantly lower than the first block (1 vs. 7: *M_diff_* = 0.029, *t*(20) = 2.40, *p* = .026, *d* = 0.52, BF_10_ = 2.28), and reached its lowest point overall in the final ten minutes of task performance (1 vs. 8: *M_diff_* = 0.062, *t*(20) = 4.23, *p* < .001, *d* = 0.92, BF_10_ = 79.2), despite equal levels of self-reported motivation. These changes in *A*′ appeared to be mostly driven by hit rate (Figure 2C), as hit rate also worsened significantly with time-on-task (*F*(2.45, 48.92) = 8.74, *p* < .001, *η*^2^ = .30), and improved transiently after the motivation manipulation (6 vs. 7: *M_diff_* = −0.077, *t*(20) = −2.17, *p* = .042, *d* = −0.47, BF_10_ = 1.56). False alarm rate appeared to decrease slightly over time (Figure 2D), but not significantly (*F*(3.08, 61.69) = 2.30, *p* = .084, *η*^2^ = .10), and was also not significantly affected by motivation (6 vs. 7: *M_diff_* = 0.006, *t*(20) = 1.10, *p* = .286, *d* = 0.24, BF_01_ = 2.59). False alarms were rare, as false alarm rate was already near the floor at the start of the experiment (3.1% of non-target trials). Finally, although response time showed the same pattern as accuracy (Figure 2B), response time did not significantly change after the motivation manipulation (6 vs. 7: *M_diff_* = 22, *t*(20) = 1.45, *p* = .163, *d* = 0.32, BF_01_ = 1.78).

### 3.2. Attentional stability: theta phase

Inter-trial phase clustering (ITPC) indexes the consistency in timing with which frequency-band-limited activity is elicited from trial to trial. Theta-band ITPC has been interpreted as a marker of attentional stability (Lutz et al., 2009). We therefore hypothesized that theta-band ITPC would decrease with time-on-task. Our data show clear ITPC in a theta-band (3–7 Hz) response from 150-500 ms post-stimulus (Figure 3A). The response was maximal over left and right parieto-occipital (lPO, rPO) electrodes, as well as over mid-frontal (MF) scalp sites (Figure 3B).

**Figure 3:**
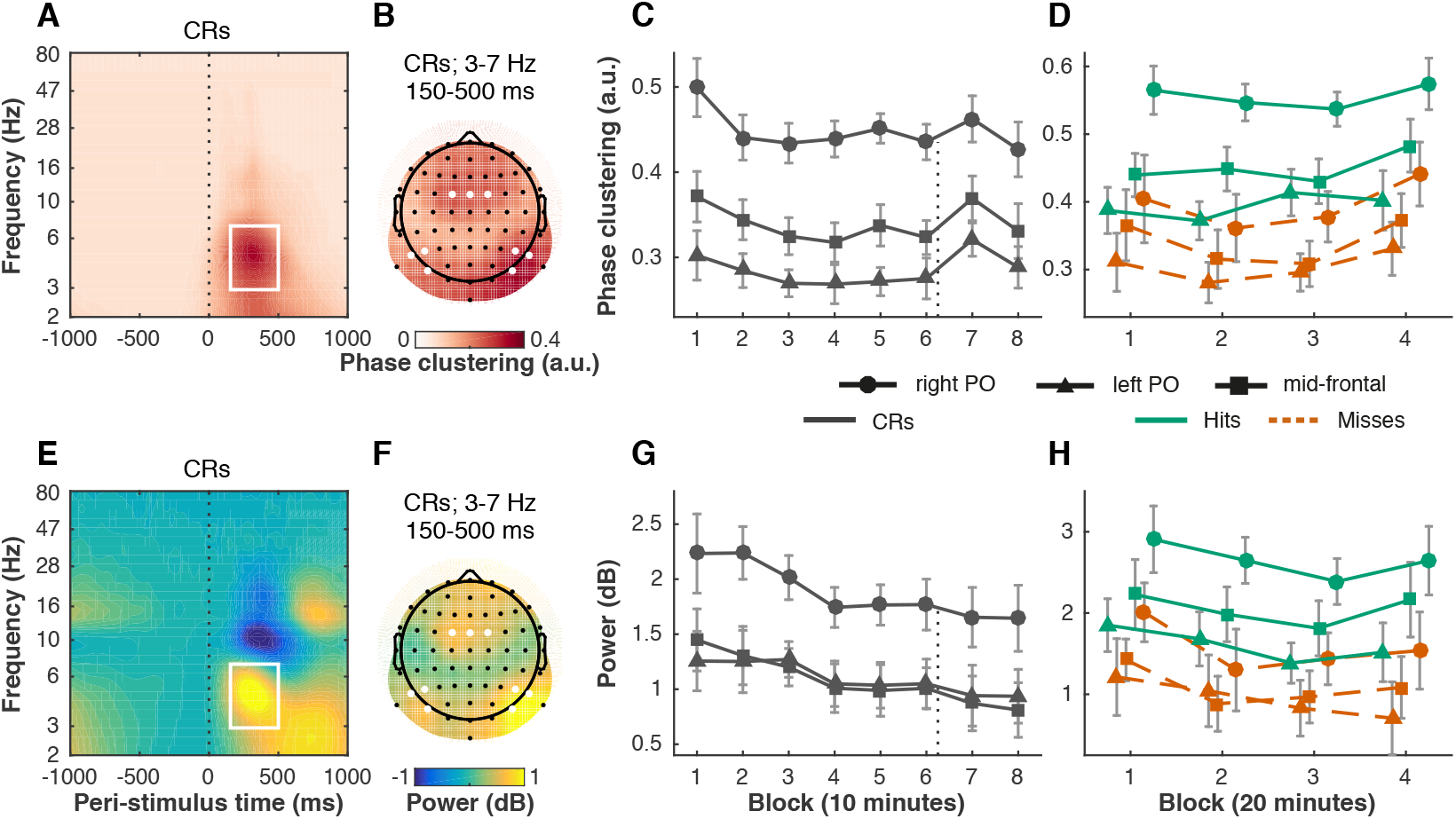
Changes in behavioral performance with time-on-task and motivation are closely tracked by cross-trial consistency of post-stimulus theta phase (A–D), but not theta power (E–H). Theta inter-trial phase clustering (ITPC) peaked from 150–500 ms post-stimulus (vertical dotted-line) between 3 and 7 Hz (time-frequency window outlined in white) (**A**, average of electrodes of interest) on left parieto-occipital (left PO) electrodes (PO7, P5, P7), right parieto-occipital (right PO) electrodes (PO8, P6, P8), and mid-frontal (MF) electrodes (FC1, FCz, FC2) (**B**, average of time-frequency window, electrodes marked in white). The same electrode sites and time-frequency windows of interest were used for theta power (**E, F**), baseline corrected from −400 to −100 ms using a decibel conversion. Theta ITPC in correct rejection trials (CRs) decreased with time-on-task (8 blocks of 10 minutes each) and increased after the motivation manipulation (directly after block 6; vertical dotted line) (**C**), closely resembling changes in task performance (Figure 2A). Power in the theta band decreased linearly over time, but did not change after the motivation manipulation (**G**). Both theta ITPC (**D**) and power (**H**) were higher for hit than miss trials, but this difference did not change significantly over time (4 blocks of 20 minutes each). Error bars are within-subject (Cousineau, 2005; Morey, 2008) 95% confidence intervals.

Over the course of the eight task blocks, theta ITPC closely tracked the time course of behavioral task performance (accuracy in *A*′) (Figure 3C). Like behavioral performance, theta ITPC decreased with time-on-task (*F*(3.83, 76.52) = 4.46, *p* = .003, 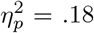) and increased directly after the motivation manipulation at two out of the three scalp regions (left PO, 6 vs. 7: *t*(20) = −3.56, *p* = .002, *d* = −0.78, BF_10_ = 20.1; right PO, 6 vs. 7: *t*(20) = −1.94, *p* = .067, *d* = −0.42, BF_10_ = 1.09; MF, 6 vs. 7: *t*(20) = −3.13, *p* = .005, *d* = −0.68, BF_10_ = 8.64).

To more directly quantify the relationship between theta ITPC and *A*′ over time, we fitted a multilevel growth model wherein we regressed the *A*′ scores on the ITPC scores in correct rejection trials. This analysis showed that ITPC significantly predicted over time (left PO: *b* = 0.11, *t*(166) = 3.11, *p* = .002; right PO: *b* = 0.12, *t*(166) = 3.89, *p* < .001; MF: *b* = 0.15, *t*(166) = 4.39, *p* < .001). ITPC was also larger in hit than in miss trials (Figure 3D, *F*(1, 20) = 58.31, *p* < .001, 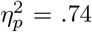), which further corroborates that the theta ITPC signal was relevant for behavior. This difference between hits and misses did not change significantly over time (no Block by Condition interaction, *F*(3, 60) = 1.39, *p* = .254, 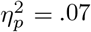).

We also investigated to what extent dynamics in theta power were similar to theta ITPC. Unsurprisingly, we observed a theta power response in the same time-frequency window and electrode sites (Figure 3E-F). Like theta ITPC, theta power decreased over time in correct rejection trials (Figure 3G, *F*(3.04, 60.85) = 4.91, *p* = .004, 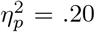). Theta power also differed between hits and misses (Figure 3H, *F*(1, 20) = 43.02, *p* < .001, 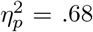). This difference did not change significantly over time (no Block by Condition interaction, *F*(3, 60) = 1.04, *p* = .383, 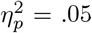). However, unlike theta ITPC, theta power was not significantly affected by the motivation manipulation (left PO, 6 vs. 7: *p* = .525, BF_01_ = 3.64; right PO, 6 vs. 7: *p* = .409, BF_01_ = 3.20; MF, 6 vs. 7: *p* = .214, BF_01_ = 2.14).

In sum, theta-band inter-trial phase clustering was higher in hit than in miss trials, suggesting it indexes behavioral performance. Changes in theta ITPC in correct rejection trials were tightly coupled to and predicted changes in behavioral performance (*A*′). Power in the theta band also decreased over time, but did not appear to respond to the motivation manipulation, suggesting that the change in power with time-on-task was partially independent of and less strongly associated with behavior than theta ITPC.

### 3.3. Early visual processing: P1 and N1 components

To investigate how time-on-task and motivation may affect early visual processing, we examined how the P1 (Figure 4A-D) and N1 (Figure 4E-H) ERP components evolved over the eight task blocks (i.e., in correct rejection trials). N1 amplitude decreased over time (Figure 4C, *F*(2.89, 57.75) = 5.79, *p* = .002, 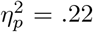), but there was no significant effect of Block for the P1 (Figure 4G, *F*(3.68, 73.70) = 1.16, *p* = .333, 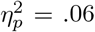). The N1 did not change significantly after the motivation manipulation, neither in the left hemisphere (6 vs. 7: *t*(20) = 0.73, *p* = .472, *d* = 0.16, BF_01_ = 3.45) nor in the right (6 vs. 7: *t*(20) = −0.33, *p* = .742, *d* = −0.07, BF_01_ = 4.18), suggesting that the decrease in the N1 with time-on-task is not sensitive to motivation, and may reflect other factors such as habituation to the repeatedly presented stimulus.

**Figure 4:**
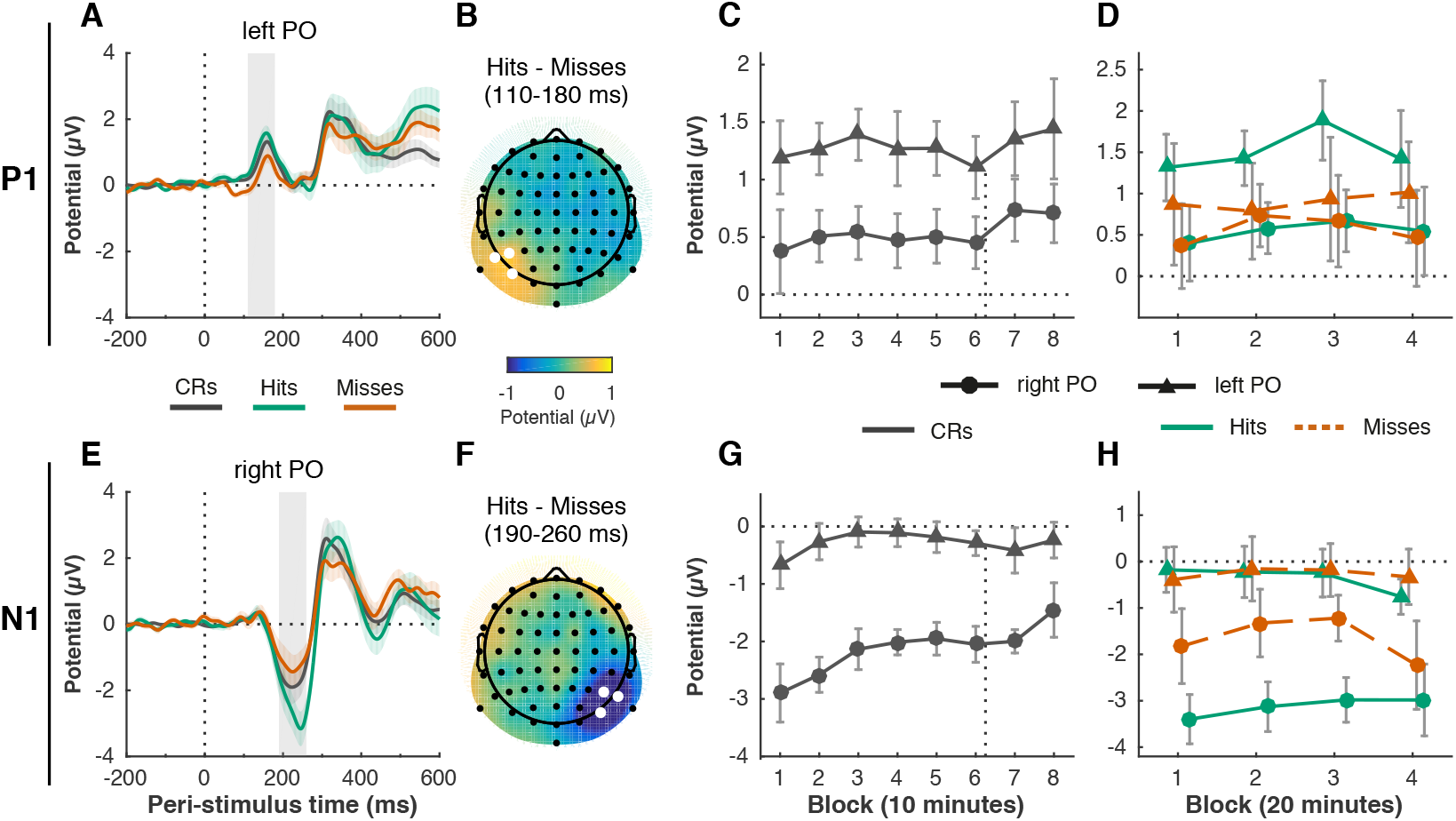
Time-on task and motivation do not affect attentional modulation of early visual processing: P1 (A-D) and N1 (E-H) ERP components. The P1 peaked between 110–180 ms (gray shaded rectangle) post-stimulus (vertical dotted line) on left parieto-occipital (left PO) electrodes (PO7, P5, P7) (**A**). The N1 peaked between 190–260 ms (gray shaded rectangle) on right parieto-occipital (right PO) electrodes (PO8, P6, P8) (**E**). The attentional modulation of the components, defined as the difference between hits and misses, was also lateralized to the left PO region for the P1 (**B**, average of P1 time window, left PO electrodes marked in white) and the right PO region for the N1 (**F**, average of N1 time window, right PO electrodes marked in white). P1 amplitude (**C**) did not change significantly over time in correct rejection (CR) trials, but N1 amplitude did decrease with time-on-task (**G**) (8 blocks of 10 minutes each). The time of the motivation manipulation (directly after block 6) is indicated with the vertical dotted line. There was no significant change in the attentional modulation (hits vs. misses) of the P1 (**D**) and N1 (**H**) over time (4 blocks of 20 minutes each). Error bars are within-subject (Cousineau, 2005; Morey, 2008) 95% confidence intervals; shaded areas in (A) and (E) represent the standard error of the mean.

The ANOVA for correct rejection trials also revealed main effects of Hemisphere, reflecting that the P1 was only visible over a left parieto-occipital scalp region (*F*(1, 20) = 13.07, *p* = .002, 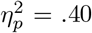), whereas the N1 peaked exclusively over the right parieto-occipital scalp region (*F*(1, 20) = 15.28, *p* < .001, 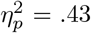). This unusual lateralization of the components is further explored elsewhere (Slagter et al., 2016). Briefly, we found that this effect could be attributed to the specific stimulus used, as well as the fact that participants were continuously attending to the left hemifield, and the right hemifield was never relevant (which differs from typical attentional cueing studies).

As top-down attention is known to increase the amplitudes of the P1 and N1, we also examined the difference between hit and miss trials, as a proxy for attentional modulation.

We hypothesized that with time on task, top-down attentional modulation of these indices of visual processing would decrease. The attentional modulations of both components were also completely lateralized (Condition by Hemisphere interaction), as both the P1 and N1 amplitude were larger for hits than misses, but only in the left and right hemisphere, respectively (P1: Figure 4B, *F*(1, 20) = 7.48, *p* = .013, 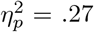; N1: Figure 4F, *F*(1, 20) = 11.86, *p* = .003, 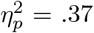). However, this effect did not interact with Block, suggesting that the attentional modulation of visual processing did not change over time or after the motivation instruction (P1: Figure 4D, *F*(3, 60) = 0.53, *p* = .664, 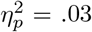; N1: Figure 4H, *F*(3, 60) = 1.70, *p* = .177, 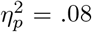).

In sum, N1 amplitude decreased with time-on-task (in correct rejection trials), but appeared to be unaffected by the motivation manipulation. The P1 and N1 components and attentional modulations thereof were completely lateralized: the P1 dissociated between hits and misses in the irrelevant hemisphere (left); the N1 dissociated between hits and misses in the relevant hemisphere (right) (see Slagter et al., 2016). However, this pattern did not change significantly with time-on-task or motivation.

### 3.4. Preparatory attentional orienting: alpha power

Next to processing of the stimulus itself, we also asked whether time-on-task might degrade preparatory attentional processes. Spatial cues that signal the location of an upcoming stimulus are known to affect the distribution of oscillatory alpha power. Specifically, alpha power decreases over the hemisphere contralateral to the attended location, and increases over the ipsilateral hemisphere. Because stimuli only appeared in the left hemifield in our task, we expected to see higher alpha power over the left (ipsilateral) hemisphere. We also expected to see this lateralization decay over time, as participants became less able to direct their attention in preparation of the stimulus.

However, both of these predictions were not borne out. First, we consistently observed higher power in a 9–14 Hz band over the right parieto-occipital scalp site than over the left (Figure 5A–B). Because we predicted exactly the opposite (higher alpha power over the left hemisphere), the alpha lateralization we observed here likely does not index top-down attentional orienting. Instead, like the unexpected lateralization in the P1 and N1 components, this inverse alpha asymmetry is probably due to our unorthodox task design where only the left hemifield was ever relevant (as further discussed in Slagter et al., 2016), and likely reflects a resting state pattern (Benwell, Keitel, Harvey, Gross, & Thut, 2018; Wieneke, Deinema, Spoelstra, Storm van Leeuwen, & Versteeg, 1980).

**Figure 5:**
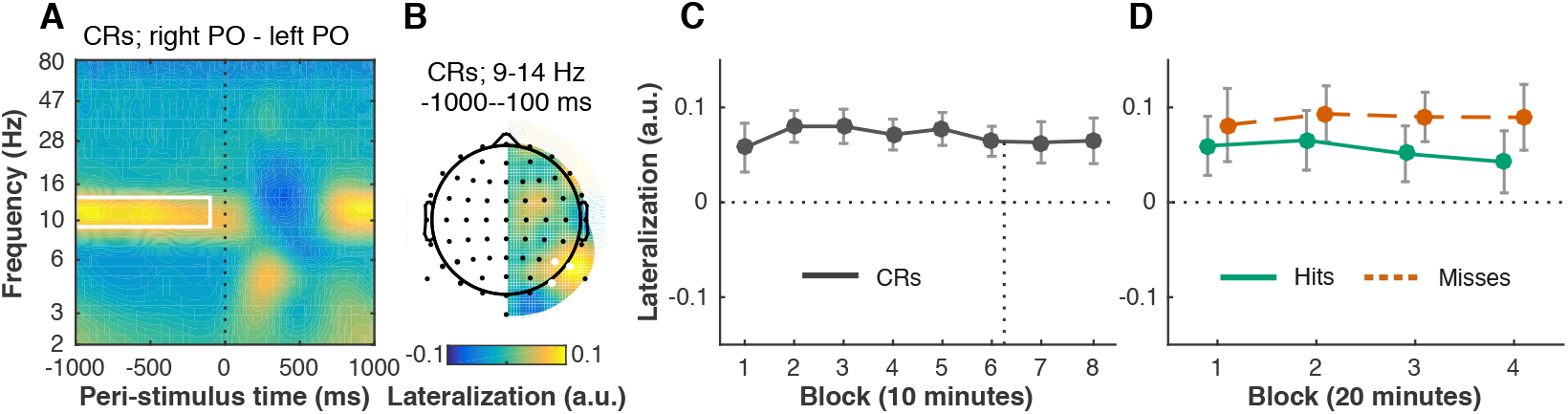
Pre-stimulus alpha power remains right-lateralized over time, despite stimuli appearing exclusively in the left visual hemifield. For each time-frequency point, we computed a lateralization index, by subtracting power at each left electrode from its counterpart on the right side, and dividing by the sum. Positive values therefore mean that power was relatively stronger over the right hemisphere. Our data show that pre-stimulus (vertical dotted line) alpha power is right-lateralized (**A**, time-frequency window: 9–14 Hz, −1000 to −100 ms, outlined in white), when comparing left (PO7, P5, P7) and right parieto-occipital (right PO: PO8, P6, P8) electrode sites (**B**, electrodes marked in white). This pattern did not change significantly with time-on-task, neither in correction rejection trials (CRs) (**C**, 8 blocks of 10 minutes each; time of the motivation manipulation indicated with vertical dotted line), nor in hit and miss trials (**D**, 4 blocks of 20 minutes each). Error bars are within-subject (Cousineau, 2005; Morey, 2008) 95% confidence intervals.

Second, this deviant pattern of alpha lateralization did not change with time-on-task (Figure 5C, *F*(3.52, 70.42) = 0.94, *p* = .438, 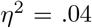, nor was it affected by the motivation manipulation (6 vs. 7: *t*(20) = 0.10, *p* = .920, *d* = 0.02, BF_01_ = 4.38). Alpha lateralization was significantly larger in miss compared to hit trials (Figure 5D, *F*(1, 20) = 12.68, *p* = .002, 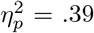) (Slagter et al., 2016), but this difference did not change significantly over time (no Block by Condition interaction, *F*(3, 60) = 0.46, *p* = .708, 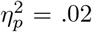.

In sum, although only one hemifield (the left) was ever relevant, we observed higher alpha power over the processing hemisphere (on the right) than the non-processing hemisphere (on the left), which is exactly opposite to the canonical pattern. This deviant alpha lateralization remained stable over time.

## 4. Discussion

We aimed to study changes in control of attention as a function of prolonged task performance, and the extent to which these could be explained by changes in motivation. Our participants performed a demanding sustained attention task for 80 minutes, and received an unexpected motivation boost after 60 minutes. We found that performance rapidly declined (i.e. the classic vigilance decrement / time-on-task effect) before the motivation boost. Afterwards, task performance did increase, but only partially (not up to the initial level) and transiently (not for the full remaining 20 minutes), even though self-reported motivation remained at initial levels.

We also recorded EEG to investigate whether changes in behavioral performance over time were accompanied by changes in three markers of attentional control. We examined pre-stimulus lateralization of alpha power as an index of preparatory orienting of attention, but did not find the canonical pattern of lateralization, nor any effect of time-on-task or motivation. In addition, the attentional modulation of the early visual P1 and N1 ERP components also did not change with time-on-task or motivation, though the absolute amplitude of the N1 decreased over time. Finally, in contrast, changes in theta-band intertrial phase clustering (ITPC) between 150–500ms post-stimulus were closely coupled to the changes in behavioral performance following time-on-task and motivation. Given that ITPC indexes the temporal consistency of neural responses across trials (VanRullen et al., 2011), the readiness of the attentional system to respond to incoming input might reduce with time-on-task. Collectively, our results thus suggest that changes in performance during sustained attention tasks are most closely associated with fluctuations in the stability of later-stage attentional processes.

### 4.1. Motivation partially and transiently restores vigilance

The sustained attention task induced a robust vigilance decrement: performance rapidly decreased with time-on-task, reaching a plateau after 20–30 minutes of task performance. After 60 minutes, we motivated our participants with an extra sum of money if they would outperform 65% of the other participants for the final 20 minutes. This prospect successfully increased self-rated levels of motivation. The motivation boost counteracted the vigilance decrement, as task performance increased in the 10-minute block immediately after participants learned of the reward. This appears inconsistent with the overload framework, which proposes that the vigilance decrement occurs due to depletion of resources—motivation alone should not be sufficient to replenish them.

However, motivation was also not enough to completely stave off the vigilance decrement, as performance was still lower than at the beginning of the task. In addition, participants appeared unable to sustain their new-found stamina: performance in the final block dropped back down to the lowest level overall. So perhaps resource depletion still played a role: participants simply could not keep up their performance, even though they were strongly incentivized to do so and self-reported to be as motivated as they were at the beginning of the task (though our Bayes factor analysis did not show strong evidence for the latter). Our findings are thus in partial agreement with both overload and motivational control accounts of the vigilance decrement.

Previous studies also found mixed evidence for the efficacy of motivation in restoring performance. In some cases, accuracy (Hopstaken et al., 2015) and response time (Boksem et al., 2006; Lorist et al., 2009) can recover after unexpected rewards, sometimes up to or beyond the initial levels. However, all of these studies measured performance at only one time point following the manipulation, so it is unknown whether performance subsequently declined—as we observed here. In contrast, two other studies have found that the slope of the vigilance decrement is not affected by low- or high reward levels (Esterman, Reagan, Liu, Turner, & DeGutis, 2014; Gergelyfi, Jacob, Olivier, & Zénon, 2015).

The main difference is that these latter two studies used trial-based rewards, which may decrease in value over time (Fortenbaugh, Degutis, & Esterman, 2017). Indeed, participants appear to discount the value of rewards when they have to pay the “cost” of sustaining their attention for longer (Massar, Lim, Sasmita, & Chee, 2016). The same goes for losses instead of rewards: when losses are small and continuous, the vigilance decrement occurs as normal. But when the risk of an instantaneous and large loss looms, the vigilance decrement can in fact be partially attenuated (Esterman et al., 2016).

These observations are in line with motivational control theories (Hockey, 1997; Kurzban et al., 2013), stating that participants are only motivated to keep up performance when benefits outweigh the costs. Note that the level of available resources may still be a principal factor in this cost/benefit computation (Boksem & Tops, 2008; Christie & Schrater, 2015). Perhaps the pattern of performance we observed—a transient increase with motivation, and a subsequent dip—also occurred because participants re-evaluated the cost/benefit ratio during the final stretches of task performance. Indeed, self-reported motivation dropped down again after the initial motivation boost, although motivation did not become significantly lower than at the start of the task.

We highlight two further caveats to our conclusion that motivation can increase performance after the initial vigilance decrement. First, we quantified performance as perceptual sensitivity (*A*′), which should exclusively reflect participants’ ability to discriminate targets from non-targets, not other factors such as response bias (the overall tendency for participants to respond ‘yes’ or ‘no’). However, Thomson, Besner, & Smilek (2016) argue that with the very low false alarm rates that vigilance tasks typically exhibit, sensitivity and response bias cannot be fully teased apart. Indeed, false alarm rate in our study was almost at floor and did not change significantly with time-on-task, so we cannot exclude that the effect on sensitivity is partially driven by response bias as well. Note that overload accounts imply a specific sensitivity decrease, but underload accounts do not necessarily.

Second, the effect that the motivation manipulation had on performance might not result from motivation per se, but due to there being a short break in the task: the 1-minute maximum period the participants had to read the instruction. Even short breaks are known to increase performance (Helton & Russell, 2015), though performance can also decrease more in subsequent task blocks after a break (Lim & Kwok, 2016). This pattern matches the changes in performance we observed after the motivation manipulation. From a strong overload perspective, one could argue that the rest afforded by the break is what caused performance to restore, and that the motivation boost played no causal role. However, the resting opportunity was very brief—all participants resumed the task within one minute— whereas these studies examined breaks of at least a minute, and much longer. Interestingly, Ross, Russell, & Helton (2014) examined the effect of 1-minute breaks in a task very similar to ours (line length discrimination). They show that breaks are effective if they occur earlier in the vigil (after 20 minutes), but not later (after 30 minutes). Because the break in our task occurred much later (after 60 minutes), the findings of Ross et al. (2014) argue against the interpretation that the subsequent increase in performance was simply due to rest.

### 4.2. Theta inter-trial phase clustering mirrors effects of time-on-task and motivation on behavioral performance

Out of our three EEG measures, post-stimulus theta ITPC was most clearly associated with prolonged performance on the sustained attention task. Theta (3–7 Hz) ITPC was larger when the target was successfully detected (hits) than when it was missed, suggesting that it indexes a behaviorally relevant process. Changes in theta ITPC closely tracked changes in behavioral performance—both those due to time-on-task as well as motivation. Theta ITPC diminished with time-on-task, which means that the timing of the neural response across trials became more variable. The increased variability might reflect that participants became progressively less able to prepare for the onset of the visual stimulus by attending at the right moment in time. We thus interpret theta ITPC as a measure of attentional stability, following previous work (Lutz et al., 2009; Slagter et al., 2009). Besides time-on-task, other factors may influence attentional stability: theta ITPC also increased after the motivation manipulation. An earlier study demonstrated that theta ITPC also decreases when participants are mind wandering (Baird, Smallwood, Lutz, & Schooler, 2014).

Oscillatory activity in the theta band is known to occur in frontal and parietal areas in visual attention tasks (Demiralp & Başar, 1992), particularly when attention has to be sustained (Clayton, Yeung, & Cohen Kadosh, 2015). The theta-band response in our study was observed over bilateral parieto-occipital electrodes, but also a mid-frontal scalp site. Of course, no conclusions on the source of this activity can be drawn based on the scalp topography, but it is possible that anterior and posterior theta-band activity reflect different processes. Frontal mid-line theta is strongly associated with cognitive control processes such as action monitoring, and likely originates from the anterior cingulate cortex (Cavanagh & Frank, 2014; Narayanan, Cavanagh, Frank, & Laubach, 2013). However, these theta-band signals are ongoing oscillations that are not phase-locked to external stimuli (Cohen & Donner, 2013), while the theta ITPC response we observe appears to be strongly phase-locked to the eliciting stimulus. Posterior theta-band activity is often observed following (any) visual stimuli (Klimesch, Sauseng, & Hanslmayr, 2007). Theta ITPC is stronger for unexpected targets, suggesting it is particularly related to attentional reorienting (Daitch et al., 2013). Theta ITPC has also been linked to matching stimuli to a memory template (Freunberger, Klimesch, Doppelmayr, & Höller, 2007; Rizzuto, Madsen, Bromfield, Schulze-Bonhage, & Kahana, 2006). Both interpretations fit our task context, as participants constantly had to evaluate whether the currently presented line matched the template of a short or long line, and targets (short lines) were more infrequent than non-targets (long lines), and thus unexpected.

Aside from the finding that theta ITPC increases with mind wandering (Baird et al., 2014), most of what is known about the relation between theta oscillations and sustained attention is about power instead of phase (Clayton et al., 2015). Several studies using different tasks have reported that frontal-midline theta power increases with time-on-task (Boksem et al., 2005; Umemoto, Inzlicht, & Holroyd, 2018; Wascher et al., 2014; but see Bonnefond et al., 2011). We also examined theta power, yet it decreased with time-on-task (over all electrodes of interest). In addition, we did not observe the same dynamics in theta power as we did in theta ITPC: power did not track behavior as closely and did not change following the motivation manipulation. So it seems that the changes in theta ITPC are independent of concurrent changes in theta power, though we cannot fully rule this out as even small differences in power can cause differences in estimation of phase (van Diepen & Mazaheri, 2018). In sum, while the exact nature of the signal remains up for debate, our results show that theta ITPC is a strong correlate of later-stage attentional or perceptual processes that are involved in the vigilance decrement.

### 4.3. No clear changes in attentional modulation of early visual P1/N1 components

It is well known that attention can modulate visual processing. Specifically, a large body of work has shown that the amplitude of the early P1 and N1 ERP components increases when a stimulus is preceded by a cue that prompts spatial attention towards it (Luck et al., 1994). As our design did not include such cues, we took the difference between hit and miss trials as a proxy for the effect of top-down attention. While P1 and N1 amplitudes were indeed larger in hit than in miss trials, this difference did not change with time-on-task. The P1 and N1 also did not respond to the motivation manipulation. We therefore conclude that neither motivation nor time-on-task affected top-down modulation of early sensory processing.

The only effect we found was a decrease in absolute N1 amplitude with time-on-task, in accord with previous studies (Boksem et al., 2005; Faber et al., 2012). However, others have previously reported that N1 amplitude remains stable with time-on-task, using tasks that more closely resemble ours (Bonnefond et al., 2010; Koelega et al., 1992). Furthermore, in the absence of any changes in the N1 attention effect, reductions in absolute N1 amplitude cannot be taken to reflect attention-related changes in visual stimulus processing, but may reflect other non-specific effects, such as habituation or perceptual learning.

### 4.4. Alpha power lateralization is reversed and unchanging

One of the most prominent EEG markers of spatial attention is the pattern of alpha lateralization: whenever one visual hemifield is attended, alpha power increases over the ipsilateral hemisphere and decreases over the contralateral hemisphere (Klimesch, 2012). In our task, all stimuli appeared left of fixation, so we expected a reduction in pre-stimulus alpha power over the right hemisphere, but instead found that alpha power was strongly right lateralized. This inverted alpha power asymmetry might be qualitatively different than the alpha lateralization that is commonly observed in spatial attention studies. In contrast to most studies, we used a task in which only the left hemifield (processed by the right hemisphere) was ever relevant, and there were no trial-to-trial attentional cues or distractors (Rihs, Michel, & Thut, 2009; Slagter et al., 2016). This particular task thus might not elicit preparatory attentional orienting on each trial. Instead, the expected stimulus location could be encoded in a more long-term form. However, it is unlikely that alpha power reflects this more sustained aspect of task knowledge instead, as alpha power was still right-lateralized when the experiment was changed such that participants sustained their attention to the right hemifield (Slagter et al., 2016) instead of the left. Either way, it is likely that alpha power does not reflect the same process in the current task context as in previous studies, so we cannot take it as an index of preparatory attentional orienting.

One earlier study found that alpha power becomes more right-lateralized with time-on-task (Newman et al., 2013). In our results, alpha power over the right hemisphere was higher compared to the left hemisphere from the start, with no change over time. A recent study by Benwell et al. (2018) reported very similar results: throughout the task (with stimuli presented at fixation), alpha power was higher over the right hemisphere than the left, but this lateralization did not change with time-on-task. Together, our results suggest that alpha power is simply right-lateralized by default, and that this resting state pattern is not susceptible to time-on-task effects. In contrast, behavioral performance typically does exhibit an increased rightward bias with time-on-task (Benwell, Harvey, Gardner, & Thut, 2013; Dufour, Touzalin, & Candas, 2007; Manly, Dobler, Dodds, & George, 2005).

Our stimuli were however always presented on the left, which could have accelerated the vigilance decrement if participants indeed orient more strongly towards the right over time. Again, because we did not observe the typical alpha lateralization pattern, we were unable to examine how spatial attentional biases might change with time-on-task.

### 4.5. Conclusion

Our study demonstrates that motivation is a key factor that may oppose the vigilance decrement, but also that motivation alone is not sufficient to fully bring task performance back online. Our results are consistent with hybrid approaches that incorporate elements from multiple frameworks (Christie & Schrater, 2015; Thomson et al., 2015), particularly motivation and resource depletion. Future studies may also include measures of mind wandering (Smallwood & Schooler, 2006) or mental fatigue (Johnston et al., 2018) to investigate how the interplay of all these factors might give rise to the vigilance decrement. We also identified that the cross-trial consistency of theta phase values—an index of attentional stability—is a close correlate of time-on-task related decreases and motivation-related increases in performance. Other EEG measures of preparatory attention (alpha power) and early visual processing (P1/N1 ERP components) were not. Larger datasets that afford more sophisticated analyses are called for to uncover how the precise relationships between different neural measures of attention and behavioral indices of vigilance unfold across time (Wang et al., 2018). At present, our findings illustrate that the vigilance decrement may not be a unitary construct, but might depend heavily on the task context and the cognitive processes that are tapped.

## Acknowledgements

We thank Katherine MacLean for sharing the task code and stimuli used in MacLean et al. (2009) with us, as well as Hilde Huizenga, Raoul Grasman and Robert Zwitser for advice on the multilevel model.

## Author contributions

**Leon Reteig**: Methodology, Software, Formal analysis, Data curation, Writing - Original draft, Visualization. **Ruud van den Brink**: Conceptualization, Methodology, Software, Investigation, Data Curation, Writing - Review & Editing, Project Administration. **Sam Prinssen**: Conceptualization, Methodology, Software, Investigation, Data Curation, Writing - Review & Editing, Project Administration. **Michael X Cohen**: Software, Resources. **Heleen Slagter**: Conceptualization, Writing - Review & Editing, Supervision, Project Administration, Funding Acquisition.

## Appendix

The following instruction was presented to participants after 60 minutes of task performance, to investigate whether motivation could improve task performance:

You have now performed this task for one hour. The last part of this experiment starts now.
During this last part, you have the opportunity to win a bonus of 30 euro’s!
This possibility is based on your task performance during the remaining 20 minutes of this experiment. Specifically, you must finish in the top 35% of participants in terms of performance.
In other words, the top-35% performers in this last part of the experiment will receive an additional 30 euro’s.
When you are certain you have seen a short line, you should respond as quickly as possible with the left mouse button.
Whenever you see a long line, do not respond.
Do your best! Good luck!

## Footnotes

3 https://doi.org/10.17605/OSF.IO/EMF9H

## References

Ackerman, P. L. (Ed.). (2011). Cognitive fatigue: Multidisciplinary perspectives on current research and future applications. Washington, DC: American Psychological Association.

Baird, B., Smallwood, J., Lutz, A., & Schooler, J. W. (2014). The decoupled mind: Mind-wandering disrupts cortical phase-locking to perceptual events. Journal of Cognitive Neuroscience, 26(11), 2596–2607. https://doi.org/10.1162/jocn_a_00656

Bartlett, F. C. (1943). Fatigue following highly skilled work. Proceedings of the Royal Society of London. Series B, Biological Sciences, 131 (864), 247–257.

Benwell, C. S., Keitel, C., Harvey, M., Gross, J., & Thut, G. (2018). Trial-by-trial co-variation of pre-stimulus EEG alpha power and visuospatial bias reflects a mixture of stochastic and deterministic effects. European Journal of Neuroscience, 48(7), 2566–2584. https://doi.org/10.1111/ejn.13688

Benwell, C. S. Y., Harvey, M., Gardner, S., & Thut, G. (2013). Stimulus- and state-dependence of systematic bias in spatial attention: Additive effects of stimulus-size and time-on-task. Cortex, 49(3), 827–836. https://doi.org/10.1016/j.cortex.2011.12.007

Boksem, M. A., Meijman, T. F., & Lorist, M. M. (2005). Effects of mental fatigue on attention: an ERP study. Cognitive Brain Research, 25(1), 107–116. https://doi.org/10.1016/j.cogbrainres.2005.04.011

Boksem, M. A., Meijman, T. F., & Lorist, M. M. (2006). Mental fatigue, motivation and action monitoring. Biological Psychology, 72(2), 123–132. https://doi.org/10.1016/j.biopsycho.2005.08.007

Boksem, M. A., & Tops, M. (2008). Mental fatigue: costs and benefits. Brain Research Reviews, 59(1), 125–139. https://doi.org/10.1016/j.brainresrev.2008.07.001

Bonnefond, A., Doignon-Camus, N., Hoeft, A., & Dufour, A. (2011). Impact of motivation on cognitive control in the context of vigilance lowering: An ERP study. Brain and Cognition, 77(3), 464–471. https://doi.org/10.1016/j.bandc.2011.08.010

Bonnefond, A., Doignon-Camus, N., Touzalin-Chretien, P., & Dufour, A. (2010). Vigilance and intrinsic maintenance of alert state: An ERP study. Behavioural Brain Research, 211 (2), 185–190. https://doi.org/10.1016/j.bbr.2010.03.030

Cajochen, C., Brunner, D. P., Krauchi, K., Graw, P., & Wirz-Justice, A. (1995). Power density in theta/alpha frequencies of the waking EEG progressively increases during sustained wakefulness. Sleep, 18(10), 890–894. https://doi.org/10.1093/sleep/18.10.890

Cavanagh, J. F., & Frank, M. J. (2014). Frontal theta as a mechanism for cognitive control. Trends in Cognitive Sciences, 18(8), 414–421. https://doi.org/10.1016/j.tics.2014.04.012

Christie, S. T., & Schrater, P. (2015). Cognitive cost as dynamic resource allocation. Frontiers in Neuroscience, 9(January), 1–23. https://doi.org/10.3389/fnins.2015.00289

Clayton, M. S., Yeung, N., & Cohen Kadosh, R. (2015). The roles of cortical oscillations in sustained attention. Trends in Cognitive Sciences, 19(4), 188–195. https://doi.org/10.1016/j.tics.2015.02.004

Cohen, M. X. (2014). Analyzing Neural Time Series Data: Theory and Practice. MIT Press.

Cohen, M. X., & Donner, T. H. (2013). Midfrontal conflict-related theta-band power reflects neural oscillations that predict behavior. Journal of Neurophysiology, 110(12), 2752–2763. https://doi.org/10.1152/jn.00479.2013

Cousineau, D. (2005). Confidence intervals in within-subject designs: A simpler solution to Loftus and Masson’s method. Tutorial in Quantitative Methods for Psychology, 1 (1), 42–45. https://doi.org/10.20982/tqmp.01.1.p042

Daitch, A. L., Sharma, M., Roland, J. L., Astafiev, S. V., Bundy, D. T., Gaona, C. M., … Corbetta, M. (2013). Frequency-specific mechanism links human brain networks for spatial attention. Proceedings of the National Academy of Sciences, 110(48), 19585–19590. https://doi.org/10.1073/pnas.1307947110

Delorme, A., & Makeig, S. (2004). EEGLAB: an open source toolbox for analysis of singletrial EEG dynamics including independent component analysis. Journal of Neuroscience Methods, 134 (1), 9–21. https://doi.org/10.1016/j.jneumeth.2003.10.009

Demiralp, T., & Başar, E. (1992). Theta rhythmicities following expected visual and auditory targets. International Journal of Psychophysiology, 13(2), 147–160.

Drapeau, C., & Carrier, J. (2004). Fluctuation of waking electroencephalogram and subjective alertness during a 25-Hour sleep-deprivation episode in young and middle-aged subjects. Sleep, 27, 55–60. https://doi.org/10.1093/sleep/27.1.55

Dufour, A., Touzalin, P., & Candas, V. (2007). Time-on-task effect in pseudoneglect. Experimental Brain Research, 176(3), 532–537. https://doi.org/10.1007/s00221-006-0810-2

Eason, R. G., Harter, M., & White, C. (1969). Effects of attention and arousal on visually evoked cortical potentials and reaction time in man. Physiology & Behavior, 4 (3), 283–289. https://doi.org/10.1016/0031-9384(69)90176-0

Eidelman-Rothman, M., Ben-Simon, E., Freche, D., Keil, A., Hendler, T., & Levit-Binnun, N. (2018). Decreased inter trial phase coherence of steady-state visual evoked responses in sleep deprivation. bioRxiv, 471730. https://doi.org/10.1101/471730

Esterman, M., Grosso, M., Liu, G., Mitko, A., Morris, R., & DeGutis, J. (2016). Anticipation of monetary reward can attenuate the vigilance decrement. PLOS ONE, 11(7), e0159741. https://doi.org/10.1371/journal.pone.0159741

Esterman, M., Noonan, S. K., Rosenberg, M., & Degutis, J. (2013). In the zone or zoning out? Tracking behavioral and neural fluctuations during sustained attention. Cerebral Cortex, 23(11), 2712–2723. https://doi.org/10.1093/cercor/bhs261

Esterman, M., Reagan, A., Liu, G., Turner, C., & DeGutis, J. (2014). Reward reveals dissociable aspects of sustained attention. Journal of Experimental Psychology: General, 143(6), 2287–2295. https://doi.org/10.1037/xge0000019

Faber, L. G., Maurits, N. M., & Lorist, M. M. (2012). Mental fatigue affects visual selective attention. PLoS ONE, 7(10), 1–10. https://doi.org/10.1371/journal.pone.0048073

Fortenbaugh, F. C., Degutis, J., & Esterman, M. (2017). Recent theoretical, neural, and clinical advances in sustained attention research. Annals of the New York Academy of Sciences, 1396, 70–91. https://doi.org/10.1111/nyas.13318

Freunberger, R., Klimesch, W., Doppelmayr, M., & Höller, Y. (2007). Visual P2 component is related to theta phase-locking. Neuroscience Letters, 426(3), 181–186. https://doi.org/10.1016/j.neulet.2007.08.062

Gergelyfi, M., Jacob, B., Olivier, E., & Zénon, A. (2015). Dissociation between mental fatigue and motivational state during prolonged mental activity. Frontiers in Behavioral Neuroscience, 9, 176. https://doi.org/10.3389/fnbeh.2015.00176

Grent-’t-Jong, T., & Woldorff, M. G. (2007). Timing and sequence of brain activity in top-down control of visual-spatial attention. PLoS Biology, 5(1), e12. https://doi.org/10.1371/journal.pbio.0050012

Händel, B. F., Haarmeier, T., & Jensen, O. (2011). Alpha oscillations correlate with the successful inhibition of unattended stimuli. Journal of Cognitive Neuroscience, 23(9), 2494–2502. https://doi.org/10.1162/jocn.2010.21557

Helton, W. S., & Russell, P. N. (2015). Rest is best: The role of rest and task interruptions on vigilance. Cognition, 134, 165–173. https://doi.org/10.106/j.cognition.2014.10.001

Helton, W. S., & Warm, J. S. (2008). Signal salience and the mindlessness theory of vigilance. Acta Psychologica, 129(1), 18–25. https://doi.org/10.1016/j.actpsy.2008.04.002

Hockey, G. R. J. (1997). Compensatory control in the regulation of human performance under stress and high workload; a cognitive-energetical framework. Biological Psychology, 45(1-3), 73–93.

Hoedlmoser, K., Griessenberger, H., Fellinger, R., Freunberger, R., Klimesch, W., Gruber, W., & Schabus, M. (2011). Event-related activity and phase locking during a psychomotor vigilance task over the course of sleep deprivation. Journal of Sleep Research, 20(3), 377–385. https://doi.org/10.1111/j.1365-2869.2010.00892.x

Hopstaken, J. F., van Der Linden, D., Bakker, A. B., & Kompier, M. A. J. (2015). A multifaceted investigation of the link between mental fatigue and task disengagement. Psychophysiology, 52(3), 305–315. https://doi.org/10.1111/psyp.12339

Johnston, D. W., Allan, J. L., Powell, D. J. H., Jones, M. C., Farquharson, B., Bell, C., & Johnston, M. (2018). Why does work cause fatigue? A real-time investigation of fatigue, and determinants of fatigue in nurses working 12-hour shifts. Annals of Behavioral Medicine, (August), 1–12. https://doi.org/10.1093/abm/kay065

Klimesch, W. (2012). Alpha band oscillations, attention, and controlled access to stored information. Trends in Cognitive Sciences, 16(12), 606–617. https://doi.org/10.1016/j.tics.2012.10.007

Klimesch, W., Sauseng, P., & Hanslmayr, S. (2007). EEG alpha oscillations: the inhibition-timing hypothesis. Brain Research Reviews, 53(1), 63–88. https://doi.org/10.1016/j.brainresrev.2006.06.003

Koelega, H. S., Verbaten, M. N., van Leeuwen, T. H., Kenemans, J. L., Kemner, C., & Sjouw, W. (1992). Time effects on event-related brain potentials and vigilance performance. Biological Psychology, 34 (1), 59–86. https://doi.org/10.1016/0301-0511(92)90024-O

Kristjansson, S. D., Kircher, J. C., & Webb, A. K. (2007). Multilevel models for repeated measures research designs in psychophysiology: An introduction to growth curve modeling. Psychophysiology, 44 (5), 728–736. https://doi.org/10.1111/j.1469-8986.2007.00544.x

Kurzban, R., Duckworth, A., Kable, J. W., & Myers, J. (2013). An opportunity cost model of subjective effort and task performance. Behavioral and Brain Sciences, 36 (06), 661–679. https://doi.org/10.1017/S0140525X12003196

Lawrence, M. A. (2016). ez: Easy Analysis and Visualization of Factorial Experiments. Retrieved from https://cran.r-project.org/package=ez

Lim, J., & Kwok, K. (2016). The effects of varying break length on attention and time on task. Human Factors, 58, 472–481. https://doi.org/10.1177/0018720815617395

Lorist, M. M., Bezdan, E., Caat, M. ten, Span, M. M., Roerdink, J. B. T. M., & Maurits, N. M. (2009). The influence of mental fatigue and motivation on neural network dynamics; an EEG coherence study. Brain Research, 1270, 95–106. https://doi.org/10.1016/j.brainres.2009.03.015

Luck, S. J., Hillyard, S. A., Mouloua, M., Woldorff, M. G., Clark, V. P., & Hawkins, H. L. (1994). Effects of spatial cueing on luminance detectability: psychophysical and electrophysiological evidence for Early selection. Journal of Experimental Psychology: Human Perception and Performance, 20(4), 887–904. https://doi.org/10.1037/0096-1523.20.4.887

Lutz, A., Slagter, H. A., Rawlings, N. B., Francis, A. D., Greischar, L. L., & Davidson, R. J. (2009). Mental training enhances attentional stability: neural and behavioral evidence. Journal of Neuroscience, 29(42), 13418–13427. https://doi.org/10.1523/JNEUROSCI.1614-09.2009

Mackworth, N. (1948). The breakdown of vigilance during prolonged visual search. Quarterly Journal of Experimental Psychology, 1, 6–21.

MacLean, K., Aichele, S., Bridwell, D., Mangun, G., Wojciulik, E., & Saron, C. (2009). Interactions between endogenous and exogenous attention during vigilance. Attention, Perception & Psychophysics, 71 (5), 1042–1058. https://doi.org/10.3758/APP.71.5.1042

Mangun, G. R., & Hillyard, S. A. (1991). Modulations of sensory-evoked brain potentials indicate changes in perceptual processing during visual-spatial priming. Journal of Experimental Psychology: Human Perception and Performance, 17(4), 1057–1074. https://doi.org/10.1037/0096-1523.17.4.1057

Manly, T., Dobler, V. B., Dodds, C. M., & George, M. A. (2005). Rightward shift in spatial awareness with declining alertness. Neuropsychologia, 43(12), 1721–1728. https://doi.org/10.1016/j.neuropsychologia.2005.02.009

Manly, T., Robertson, I., & Galloway, M. (1999). The absent mind: further investigations of sustained attention to response. Neuropsychologia, 37, 661–670. https://doi.org/10.1016/S0028-3932(98)00127-4

Massar, S. A., Lim, J., Sasmita, K., & Chee, M. W. (2016). Rewards boost sustained attention through higher effort: A value-based decision making approach. Biological Psychology, 120, 21–27. https://doi.org/10.1016/j.biopsycho.2016.07.019

Mazaheri, A., van Schouwenburg, M. R., Dimitrijevic, A., Denys, D., Cools, R., & Jensen, O. (2014). Region-specific modulations in oscillatory alpha activity serve to facilitate processing in the visual and auditory modalities. NeuroImage, 87, 356–362. https://doi.org/10.1016/j.neuroimage.2013.10.052

Morey, R. (2008). Confidence intervals from normalized data: A correction to Cousineau (2005). Tutorial in Quantitative Methods for Psychology, 4 (2), 61–64. https://doi.org/10.20982/tqmp.04.2.p061

Morey, R. D., & Rouder, J. N. (2018). BayesFactor: Computation of Bayes Factors for Common Designs. Retrieved from https://cran.r-project.org/package=BayesFactor

Mrazek, M. D., Smallwood, J., Franklin, M. S., Chin, J. M., Baird, B., & Schooler, J. W. (2012). The role of mind-wandering in measurements of general aptitude. Journal of Experimental Psychology: General, 141 (4), 788–798. https://doi.org/10.1037/a0027968

Narayanan, N. S., Cavanagh, J. F., Frank, M. J., & Laubach, M. (2013). Common medial frontal mechanisms of adaptive control in humans and rodents. Nature Neuroscience, 16(12), 1888–1895. https://doi.org/10.1038/nn.3549

Newman, D. P., O’Connell, R. G., & Bellgrove, M. A. (2013). Linking time-on-task, spatial bias and hemispheric activation asymmetry: A neural correlate of rightward attention drift. Neuropsychologia, 51 (7), 1215–1223. https://doi.org/10.1016/j.neuropsychologia.2013.03.027

O’Connell, R. G., Dockree, P. M., Robertson, I. H., Bellgrove, M. A., Foxe, J. J., & Kelly, S. P. (2009). Uncovering the neural signature of lapsing attention: electrophysiological signals predict errors up to 20 s before they occur. Journal of Neuroscience, 29(26), 8604–8611. https://doi.org/10.1523/JNEUROSCI.5967-08.2009

Parasuraman, R. (1979). Memory load and event rate control sensitivity decrements in sustained attention. Science, 205(4409), 924–927. https://doi.org/10.1126/science.472714

Pinheiro, J., Bates, D., & R-core. (2018). nlme: Linear and Nonlinear Mixed Effects Models. Retrieved from https://cran.r-project.org/package=nlme

R Core Team. (2017). R: A Language and Environment for Statistical Computing. Vienna, Austria: R Foundation for Statistical Computing. Retrieved from https://www.r-project.org/

Reteig, L. C., van den Brink, R. L., Prinssen, S., Cohen, M. X., & Slagter, H. A. (2018). Sustaining attention for a prolonged period of time increases temporal variability in cortical responses. https://doi.org/10.17605/OSF.IO/456HE

Rihs, T. A., Michel, C. M., & Thut, G. (2009). A bias for posterior alpha-band power suppression versus enhancement during shifting versus maintenance of spatial attention. NeuroImage, 44 (1), 190–199. https://doi.org/10.1016/j.neuroimage.2008.08.022

Rizzuto, D. S., Madsen, J. R., Bromfield, E. B., Schulze-Bonhage, A., & Kahana, M. J. (2006). Human neocortical oscillations exhibit theta phase differences between encoding and retrieval. NeuroImage, 31 (3), 1352–1358. https://doi.org/10.1016/j.neuroimage.2006.01.009

Robertson, I. H., Manly, T., Andrade, J., Baddeley, B. T., & Yiend, J. (1997). ‘Oops!’: performance correlates of everyday attentional failures in traumatic brain injured and normal subjects. Neuropsychologia, 35(6), 747–758. https://doi.org/10.1016/S0028-3932(97)00015-8

Robinson, D., & Hayes, A. (2018). broom: Convert Statistical Analysis Objects into Tidy Tibbles. Retrieved from https://cran.r-project.org/package=broom

Ross, H. A., Russell, P. N., & Helton, W. S. (2014). Effects of breaks and goal switches on the vigilance decrement. Experimental Brain Research, 232(6), 1729–1737. https://doi.org/10.1007/s00221-014-3865-5

Rouder, J. N., Speckman, P. L., Sun, D., Morey, R. D., & Iverson, G. (2009). Bayesian t tests for accepting and rejecting the null hypothesis. Psychonomic Bulletin and Review, 16(2), 225–237. https://doi.org/10.3758/PBR.16.2.225

R Studio Team. (2016). RStudio: Integrated Development for R. Boston, MA: RStudio, Inc.

Sauseng, P., Klimesch, W., Stadler, W., Schabus, M., Doppelmayr, M., Hanslmayr, S., … Birbaumer, N. (2005). A shift of visual spatial attention is selectively associated with human EEG alpha activity. The European Journal of Neuroscience, 22(11), 2917–2926. https://doi.org/10.1111/j.1460-9568.2005.04482.x

Seli, P., Cheyne, J. A., Xu, M., Purdon, C., & Smilek, D. (2015). Motivation, intentionality, and mind wandering: Implications for assessments of task-unrelated thought. Journal of Experimental Psychology: Learning, Memory, and Cognition, 41 (5), 1417–1425. https://doi.org/10.1037/xlm0000116

Seli, P., Schacter, D. L., Risko, E. F., & Smilek, D. (2017). Increasing participant motivation reduces rates of intentional and unintentional mind wandering. Psychological Research, 1–13. https://doi.org/10.1007/s00426-017-0914-2

Simmons, J. P., Nelson, L. D., & Simonsohn, U. (2012). A 21 Word Solution. Dialogue, the Official Newsletter of the Society for Personality and Social Psychology, 1–4. https://doi.org/10.2139/ssrn.2160588

Slagter, H. A., Lutz, A., Greischar, L. L., Nieuwenhuis, S., & Davidson, R. J. (2009). Theta phase synchrony and conscious target perception: impact of intensive mental training. Journal of Cognitive Neuroscience, 21 (8), 1536–1549. https://doi.org/10.1162/jocn.2009.21125

Slagter, H. A., Prinssen, S., Reteig, L. C., & Mazaheri, A. (2016). Facilitation and inhibition in attention: Functional dissociation of pre-stimulus alpha activity, P1, and N1 components. NeuroImage, 125, 25–35. https://doi.org/10.1016/j.neuroimage.2015.09.058

Smallwood, J., & Schooler, J. W. (2006). The restless mind. Psychological Bulletin, 132(6), 946–958. https://doi.org/10.1037/0033-2909.132.6.946

Stanislaw, H., & Todorov, N. (1999). Calculation of signal detection theory measures. Behavior Research Methods, Instruments, & Computers, 31 (1), 137–149.

Talsma, D., Mulckhuyse, M., Slagter, H. A., & Theeuwes, J. (2007). Faster, more intense! The relation between electrophysiological reflections of attentional orienting, sensory gain control, and speed of responding. Brain Research, 1178(1), 92–105. https://doi.org/10.1016/j.brainres.2007.07.099

Taylor, M., & Creelman, C. (1967). PEST: Efficient Estimates on Probability Functions. The Journal of the Acoustic Society of America, 41 (4), 782–787.

Thomson, D. R., Besner, D., & Smilek, D. (2015). A Resource-Control Account of Sustained Attention: Evidence From Mind-Wandering and Vigilance Paradigms. Perspectives on Psychological Science, 10(1), 82–96. https://doi.org/10.1177/1745691614556681

Thomson, D. R., Besner, D., & Smilek, D. (2016). A critical examination of the evidence for sensitivity loss in modern vigilance tasks. Psychological Review, 123(1), 70–83. https://doi.org/10.1037/rev0000021

Thut, G., Nietzel, A., Brandt, S. A., & Pascual-Leone, A. (2006). Alpha-band electroencephalographic activity over occipital cortex indexes visuospatial attention bias and predicts visual target detection. Journal of Neuroscience, 26(37), 9494–9502. https://doi.org/10.1523/JNEUROSCI.0875-06.2006

Umemoto, A., Inzlicht, M., & Holroyd, C. B. (2018). Electrophysiological indices of anterior cingulate cortex function reveal changing levels of cognitive effort and reward valuation that sustain task performance. Neuropsychologia. https://doi.org/10.1016/j.neuropsychologia.2018.06.010

van den Brink, R. L., Murphy, P. R., & Nieuwenhuis, S. (2016). Pupil diameter tracks lapses of attention. PLoS ONE, 11 (10), 1–16. https://doi.org/10.1371/journal.pone.0165274

van Diepen, R. M., & Mazaheri, A. (2018). The caveats of observing inter-trial phase-coherence in cognitive neuroscience. Scientific Reports, 8(1), 1–9. https://doi.org/10.1038/s41598-018-20423-z

VanRullen, R., Busch, N. A., Drewes, J., & Dubois, J. (2011). Ongoing EEG phase as a trial-by-trial predictor of perceptual and attentional variability. Frontiers in Psychology, 2, 60. https://doi.org/10.3389/fpsyg.2011.00060

Wang, H.-T., Smallwood, J., Mourao-Miranda, J., Xia, C. H., Satterthwaite, T. D., Bassett, D. S., & Bzdok, D. (2018). Finding the needle in high-dimensional haystack: A tutorial on canonical correlation analysis. arXiv, 1812.02598. Retrieved from http://arxiv.org/abs/1812.02598

Warm, J. S., Parasuraman, R., & Matthews, G. (2008). Vigilance Requires Hard Mental Work and Is Stressful. Human Factors, 50(3), 433–441. https://doi.org/10.1518/001872008X312152

Wascher, E., Rasch, B., Sänger, J., Hoffmann, S., Schneider, D., Rinkenauer, G., … Gutberlet, I. (2014). Frontal theta activity reflects distinct aspects of mental fatigue. Biological Psychology, 96(1), 57–65. https://doi.org/10.1016/j.biopsycho.2013.11.010

Wickham, H. (2017). tidyverse: Easily Install and Load the ‘Tidyverse’. Retrieved from https://cran.r-project.org/package=tidyverse

Wieneke, G. H., Deinema, C. H. A., Spoelstra, P., Storm van Leeuwen, W., & Versteeg, H. (1980). Normative spectral data on alpha rhythm in male adults. Electroencephalography and Clinical Neurophysiology, 49(5-6), 636–645. https://doi.org/10.1016/0013-4694(80)90404-6

Worden, M. S., Foxe, J. J., Wang, N., & Simpson, G. V. (2000). Anticipatory biasing of visuospatial attention indexed by retinotopically specific alpha-band electroencephalography increases over occipital cortex. Journal of Neuroscience, 20(6), 1–6. https://doi.org/10.1523/JNEUROSCI.20-06-j0002.2000

Xie, Y. (2015). Dynamic Documents with R and knitr. (J. M. Chambers, T. Hothorn, D. T. Lang, & H. Wickham, Eds.) (2nd editio). Boca Raton, FL, USA: CRC Press.

Xie, Y. (2018). knitr: A General-Purpose Package for Dynamic Report Generation in R. Retrieved from https://cran.r-project.org/package=knitr

